# GraphTME: a framework for predicting response to immune checkpoint inhibitors by interpreting cell-cell interactions in the tumour microenvironment using spatial transcriptomics of tumour tissue

**DOI:** 10.1101/2025.07.24.665835

**Authors:** Hoyeon Jeong, Junghan Oh, Yoon-La Choi

## Abstract

Immune checkpoint inhibitors (ICIs) have been used to treat cancer by reactivating T-cell responses against tumours, yet clinical efficacy remains limited due to low response rates and the lack of robust predictive biomarkers. The tumour microenvironment critically shapes ICI responsiveness by modulating antitumour immunity, necessitating a deeper understanding of spatially organised cell–cell interactions. Imaging-based spatial transcriptomics (ST) enables such analysis at single-cell resolution. We present GraphTME, a spatially informed and biologically interpretable framework that models pathway-specific directional cell–cell interactions to predict anti-PD-1 response. Ligand–receptor interactions are organised by signalling pathways and represented as a multi-relational directed graph, with edge weights inversely scaled by spatial distance. Using CD8^+^ T cells from single-cell RNA-seq data of ICI-treated NSCLC patients, we trained a relational graph convolutional network to infer immune responsiveness. GraphTME achieved an *F*_1_ score exceeding 0.83 in predicting ICI response and was validated using MERFISH data from an NSCLC patient with a partial response. Predicted responder CD8^+^ T cells exhibited higher abundance and stronger directional signalling to tumour cells. They also expressed genes associated with antitumour activity, while non-responders showed expression patterns linked to poor prognosis. GraphTME is the first framework to quantitatively model single-cell-level interactions within the spatial architecture of tumour tissues and predict ICI responses from these interactions. It offers a spatially resolved, biologically grounded biomarker for immunotherapy and a tool for dissecting immune dynamics *in situ*.

## Introduction

Immune checkpoint inhibitors (ICIs) have emerged as key modalities in cancer immunotherapy by reactivating cytotoxic T lymphocytes, particularly CD8^+^ T cells, against tumour cells [1]. This is achieved by blocking immune checkpoints, such as PD-1/PD-L1 and CTLA-4, thereby enhancing the recognition of T cell receptors and CD8 coreceptors in peptide–MHC complexes present on the surface of cancer cells. However, despite their success in certain patient populations, ICIs have limited response rates, highlighting the need for precise biomarkers to guide treatment [2]. Currently, PD-L1 immunohistochemistry is widely used in the clinical setting to stratify patients likely to respond to ICIs. In parallel, next-generation sequencing-based assessments such as tumour mutational burden and neoantigen prediction have been investigated as complementary biomarkers [3, 4]. Nonetheless, these approaches largely focus on tumour-intrinsic features and often fail to capture the complexity of the tumour microenvironment (TME), which plays a critical role in modulating therapeutic response. The TME is largely composed of vascular endothelial cells that promote cancer cell growth and suppress T cells, and it is important to understand the interactions between these constituent cells for effective cancer treatment [5]. Understanding these spatially organised intercellular interactions is essential for identifying the determinants of ICI responsiveness beyond the intrinsic properties of cancer cells [6].

Spatial transcriptomics (ST) is a useful platform for capturing gene expression data while preserving the spatial context of tissue sections. Imaging-based ST technologies that use fluorescence *in situ* hybridisation, such as MERFISH and Xenium, provide subcellular resolution and single-cell segmentation, making them particularly suitable for analysing cell–cell interactions within the TME [7, 8]. For instance, a recent study using MERSCOPE observed that fibroblasts recruit T cells and link them to the ICI response in non-small cell lung cancer (NSCLC) [9]. However, that study lacked a quantitative modelling framework and reproducible prediction capability, limiting its translational impact.

Recent advances in ST have spurred the development of computational frameworks to model cell-cell interactions (CCIs) within the TME. Several representative models have been proposed to leverage spatial and molecular contexts for interaction inference and therapeutic response prediction. HoloNet [10], SpaCI [11], and SpaRx [12] use graph-attention-based mechanisms. These models primarily focus on CCIs between predefined cell types. The SpaCI study predicted immune patterns by simulating ligand-receptor interactions in MERSCOPE data from patients with colon cancer and melanoma using adaptive graph attention. Spacia [13] extended the modelling scope to include both intra- and intercellular interactions using a Bayesian multiple-instance learning framework. This study used MERSCOPE data from patients with prostate cancer to infer interactions between cells in the TME, but the inferred interactions were not visualised or quantified in the context of tumour-immune modulation. SpaTalk [14] aims to resolve CCIs at a single-cell resolution using a biological knowledge graph. SpaCET [15] also applies spatial colocalization at single-cell resolution via Spearman correlation, demonstrating a tumour-immune interface, particularly between cancer-associated fibroblasts and M2 macrophages. Nonetheless, none of the models quantify spatial interactions at the level of individual cells within the TME or addressed the prediction of immunotherapy responses. Therefore, in this study, we developed GraphTME, a relational graph convolutional network (RGCN) [16]-based model for ligand-receptor interactions between individual cells based on spatial proximity within the TME cells, to quantitatively predict ICI responses.

## Materials and methods

The GraphTME framework for ICI response prediction comprises two phases: (1) a training phase using single-cell RNA-seq data from CD8^+^ T cells of patients with NSCLC treated with anti-PD-1 therapy (Fig. 1A), and (2) an inference phase in which the trained GraphTME model is applied to ST data with characterised cell–cell interactions to predict individual T cell responsiveness *in situ* (Fig. 1B).

**Fig. 1:**
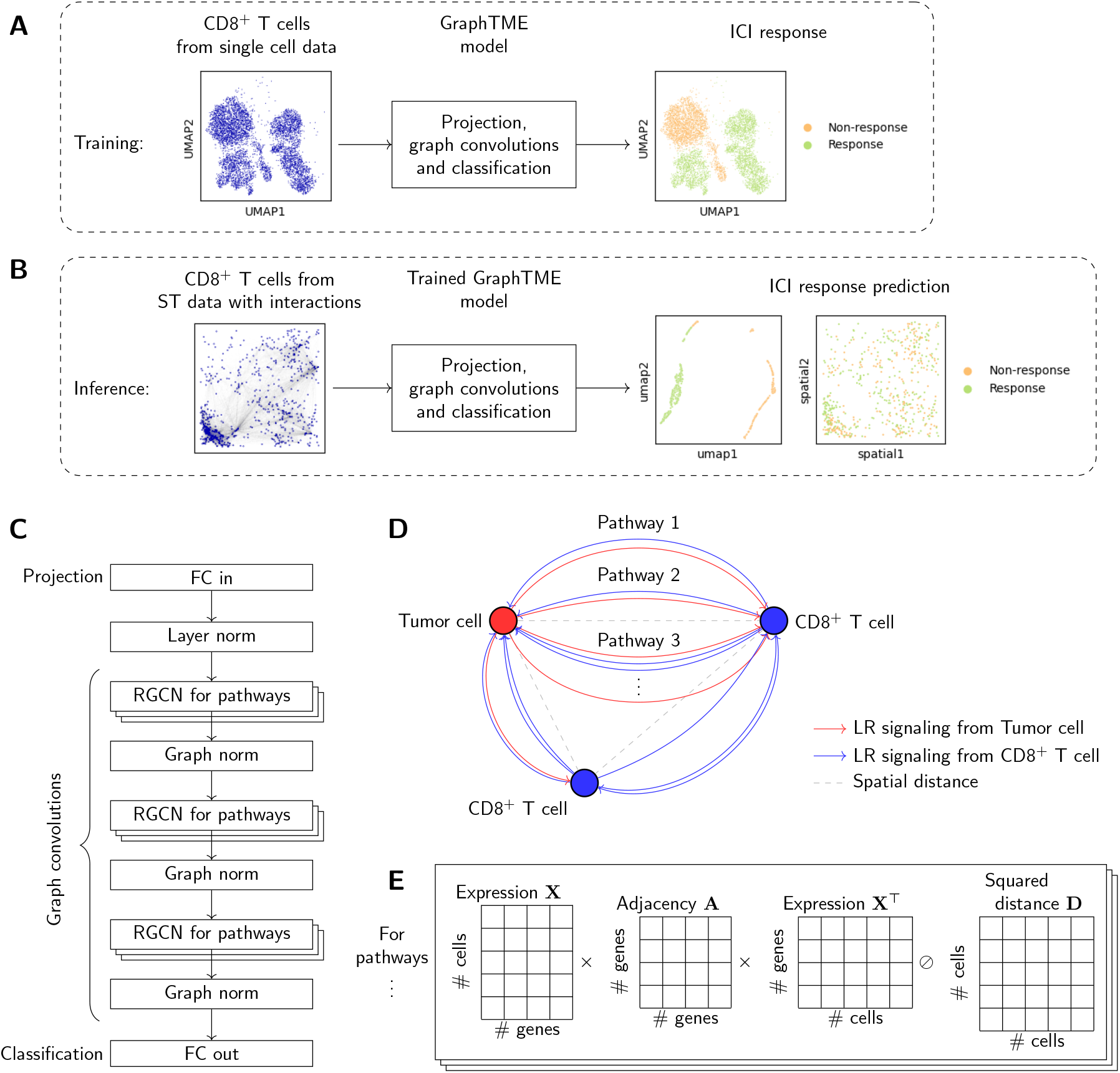
Schematic of an ICI response prediction model based on cell–cell interaction (CCI) graphs. **A**. Training of GraphTME models using the CD8^+^ T cells from single-cell data; **B**. Inference using CD8^+^ T cells from spatial transcriptomics data at single-cell resolution with the trained GraphTME model; **C**. GraphTME model architecture, consisting of an input projection layer for dimensionality reduction, extraction layers for node features and edge features by pathway, and a classifier layer; **D**. Schematic diagram of CCIs, showing mutual distances and directed edges between ligands and receptors along pathways; **E**. Adjacency matrix operation of CCIs by pathways: converts gene-level ligand-receptor interactions into a weighted matrix between cells based on expression levels and makes it inversely proportional to intercellular distance.

### Database for a pipeline of cell-level analysis

To construct various steps of a computational pipeline for cell-level analysis—including cell-type annotation, ligand-receptor interaction modelling, and ICI response prediction—we used publicly available datasets curated from established resources. Table 1 summarises these datasets and their applications.

**Table 1:** External databases used in the analysis pipeline.

| Usage | Data source | Data type | Cancer type used | Cell types used |
| --- | --- | --- | --- | --- |
| Modelling ICI response prediction | Kagamu et al. [17] | Chromium | NSCLC | CD8<sup>+</sup> T cells |
| Reference for cell type annotation | CZ CELLxGENE Discover | Various single-cell RNA-seq | LUAD, LUSC, NSCLC and breast cancer | All major TME cell types |
| Linking ligand-receptor interactions | TICCom | Gene symbol pairs | LUAD, NSCLC and breast cancer | N/A |

Abbreviations: ICI, immune checkpoint inhibitor; LUAD, lung adenocarcinoma; LUSC, lung squamous cell carcinoma; NSCLC, non-small cell lung cancer; TME, tumour microenvironment.

To train the ICI response prediction model, we used CD8^+^ T cells extracted from the Chromium-based single-cell RNA-seq dataset presented by Kagamu et al [17]. (Fig. 1A). This dataset contains clinical annotations based on the RECIST 1.1 criteria [18], enabling supervised learning of response and non-response profiles.

For cell type annotation, we used single-cell RNA-seq reference data from the CZ CELLxGENE Discover Portal [19]. These references, encompassing diverse cell populations from lung and breast cancer tissues, were used to label cells in ST.

To model CCIs, we leveraged the TICCom database [20], which organises ligand-receptor signalling between TME cell types into pathways, from which directed edges can be constructed to build pathway-specific interaction graphs.

### Construction of the GraphTME model based on relational graph convolutional networks

To predict the ICI response from CD8^+^ T cells, we constructed a relational graph convolutional network (RGCN) [16] to model the pathway-specific signalling relationships between cells. The Python code was implemented using the Pytorch Geometric library [21]. The model operates on a multi-relational graph 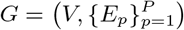, where *V* = {*v*_1_, …, *v*_*n*_} is the set of *n* nodes, each corresponding to a CD8^+^ T cell, and *E*_*p*_ denotes the set of directed edges of pathway type *p*, each corresponding to a ligand-receptor interaction in pathway *p* ∈ {1, …, *P*}. Edges are defined as (*i, j, w*) for all *E*_*p*_, indicating a directed connection from node *v*_*i*_ to node *v*_*j*_ with weight *w* > 0 under relation type *p*. During training, the graph contained only node features derived from transcriptomic profiles and no edge connectivity. Thus, the input training graph is defined as *G*_train_ = (*V*, ∅). During inference, we used multi-relational edges *E*_*p*_ derived from pathway-specific ligand-receptor interactions.

The GraphTME model consists of three main components (Fig. 1C). (1) The input projection layer applies a linear transformation **X**^(1)^ = relu **XW**^(0)^, mapping the input features to a hidden space. Here **X** is the node feature matrix defined as **X** ∈ ℝ^*n*×*g*^, where *n* is the number of CD8^+^ T cells, and *g* is the dimensionality of the gene expression feature space. Subsequently, layer normalisation [22] and dropout (rate 0.3) are applied. **W**^(0)^ is the trainable weight matrix in the first layer, and relu denotes the ReLU activation function. (2) The hidden representation stage consists of three stacked RGCN layers. Each layer *l* applies relational aggregation and transformation as 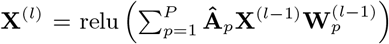, where **Â**_*p*_ is the normalised adjacency matrix for pathway *p*, and 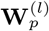 is the trainable weight matrix for pathway *p* in layer *l*. Graph normalisation [23] and dropout (rate 0.3) are applied after each layer. (3) The output layer is a fully connected linear layer **Ŷ** = softmax **X**^(*L*−1)^**W**^(*L*−1)^, projecting onto a two-class softmax space for ICI response classification (response or non-response).

### Training the model

The model was trained in a supervised manner using single-cell RNA-seq data from patients with NSCLC treated with ICI (Fig. 1A). CD8^+^ T cells were extracted from each sample and annotated with binary labels derived from RECIST 1.1: response (CR and PR) or non-response (SD and PD). To correct for technical and batch effects, latent embedding extraction was performed using the scVI framework [24]. The resulting features were used as inputs for the RGCN.

The model was optimised using the cross-entropy loss ℒ = − Σ_*i*_ *y*_*i*_ log *ŷ*_*i*_, where *y*_*i*_ ∈ {0, 1} is the ground truth label and *ŷ*_*i*_ is the model-predicted probability. Training was performed over 400 epochs using the Adam optimiser with a learning rate of 0.01. To ensure reproducibility, all random seeds were fixed across the NumPy [25], PyTorch, and CUDA backends. The training and inference were conducted on an NVIDIA H100 GPU with CUDA acceleration.

### ST datasets acquisition

ST datasets were obtained from multiple platforms and sources, covering both lung and breast cancer samples. The datasets were collected from public repositories and published literature, as detailed in Table 2. Xenium Prime datasets were acquired from the 10X Genomics public data portal and included lung and breast cancer tissues. The MERFISH dataset from Chen et al. [9] included an NSCLC sample with a known clinical response to pembrolizumab (anti-PD-1 therapy). This sample was annotated as a partial response (PR) based on RECIST 1.1, providing an opportunity to validate the predictive modelling of ICI outcomes.

**Table 2:**
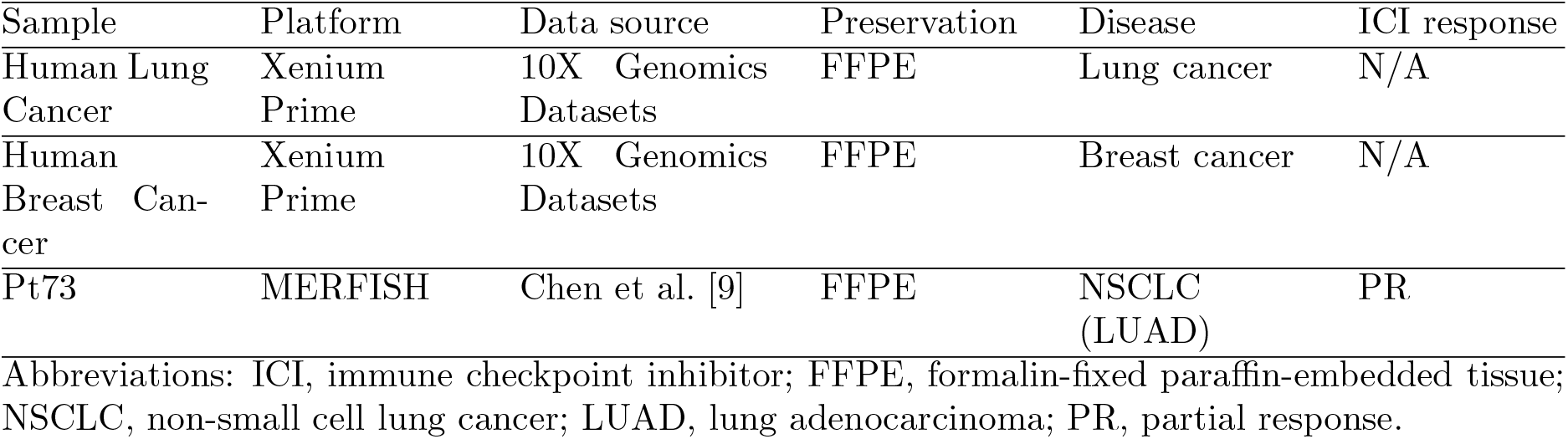
Information for each sample across platforms.

| Sample | Platform | Data source | Preservation | Disease | ICI response |
| --- | --- | --- | --- | --- | --- |
| Human Lung Cancer | Xenium Prime | 10X Genomics Datasets | FFPE | Lung cancer | N/A |
| Human Breast Cancer | Xenium Prime | 10X Genomics Datasets | FFPE | Breast cancer | N/A |
| Pt73 | MERFISH | Chen et al. [9] | FFPE | NSCLC (LUAD) | PR |

Abbreviations: ICI, immune checkpoint inhibitor; FFPE, formalin-fixed paraffin-embedded tissue; NSCLC, non-small cell lung cancer; LUAD, lung adenocarcinoma; PR, partial response.

### Data preprocessing and cell type annotation

The ST data were processed using Scanpy [26]. Mitochondrial quality control filtering was performed to remove low-quality cells. To enable comparison across gene groups, expression values were normalised to 10,000 and logarithmised. Cell types were annotated using the TACCO framework [27], leveraging single-cell RNA-seq reference atlases specific to the cancer type.

### Selection of a representative spatial region

To prioritise biologically meaningful regions for downstream CCI analysis, we developed a grid-based spatial screening strategy to identify the most representative tumour–immune interface. This method systematically searches for ST tissue sections in regions characterised by both immune infiltration and spatial heterogeneity.

Let Ω denote the spatial domain of a tissue section and define a partition 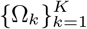 of Ω into non-overlapping 1 mm × 1 mm square regions. For each region Ω_*k*_, we computed the following three key metrics: (1) 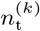 is the number of CD8^+^ T cells in region Ω_*k*_; (2) 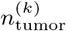 is the number of tumour cells in region Ω_*k*_; and (3) 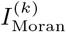 is Moran’s I statistic for the spatial distribution of two cell types in Ω_*k*_. The region-wise selection score is defined as 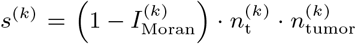. This formulation ensures that the score *s* increases with the abundance of both immune and tumour cells, while also favouring regions with lower spatial autocorrelation. In particular, lower values of Moran’s *I* indicate reduced spatial clustering and greater intermixing between cell types, which may reflect more active tumour–immune interactions. The PySAL library [28] was used for spatial autocorrelation analysis. Finally, the region 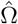 selected for analysis is 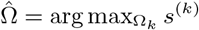.

### Graph construction with directed ligand-receptor interaction for each pathway

To model signalling-based CCIs, we constructed pathway-specific graphs using ligand-receptor interaction data from the TICCom database (Fig. 1D). For each curated signalling pathway, we generated a sparse, directed adjacency matrix representing gene-gene connectivity and then propagated gene-level information through cell-level expression profiles. Fig. 1E shows the distance-normalised interaction matrix between cells. Linear algebraic computations for the graph construction were performed using the NumPy library. Each directed edge from a source cell to a target cell was quantified by ligand-receptor co-expression, scaled by spatial distance-adjusted normalisation. Let **X** ∈ ℝ^*N*×*g*^ be the cell-by-gene expression matrix, where *N* is the number of cells across all cell types, and *g* is the number of genes. Let 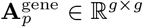 denote the directed ligand-receptor adjacency matrix for pathway *p*, where the (*i, j*)th entry encodes the interaction strength from ligand *i* to receptor *j*. We compute a CCI matrix 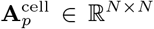 for each pathway *p* as 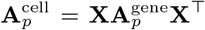. Here, 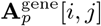 quantifies the potential signalling from cell *i* to cell *j* via pathway *p* by aggregating ligand expression in cell *i* and receptor expression in cell *j* using known ligand-receptor pairs. To incorporate spatial proximity, we compute a pairwise distance matrix **D** ∈ ℝ^*N*×*N*^, where **D**[*i, j*] is the squared Euclidean distance between cells *i* and *j* based on their spatial coordinates. The distance-normalised interaction strength is given by

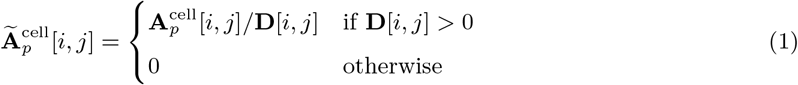

This penalises long-range interactions, ensuring that interactions between spatially distant cells are downweighted. From the resulting matrix 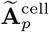, we extract all nonzero entries as weighted directed edges, forming the edge list *E*_*p*_ = {(*i, j, w*_*ij*_) | **Ã**_*p*_[*i, j*]^cell^ > 0}. Repeating this process for each pathway *p* yields a collection of graphs *G*_*p*_ = (*V, E*_*p*_), where *V* is the set of all cells and *E*_*p*_ encodes the pathway-specific directed interactions. These graphs collectively form a multi-relational graph structure 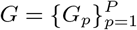, which serves as the input to the RGCN model.

### Inference of ICI responses to the ST data

To validate anti-PD-1 ICI responsiveness in the ST data, we performed inference on CD8^+^ T-cells located within the previously selected 1 mm^2^ representative regions (Fig. 1B). Using the trained GraphTME model, we predicted single-cell-level response probabilities. Following inference, CD8^+^ T cells were spatially projected and classified as responders and non-responders. First, we compared the feature distributions of responder and non-responder T cells by projecting the node embeddings from the final RGCN layer into a two-dimensional latent space using UMAP. Second, we overlaid the predicted labels onto the original tissue coordinates to visualise response heterogeneity across the tumour microenvironment.

### Data representation and visualisation

Visualisations of the results were produced using widely adopted, peer-reviewed Python libraries to ensure clarity, reproducibility. Multi-framed layouts were arranged using Matplotlib [29]. Spatial maps, UMAP projections and dot plots of gene expression were rendered with Scanpy, while bar plots and heatmaps were created using Seaborn [30]. Graph-structured data, including cell–cell interaction networks, were visualised with NetworkX [31]. Directed signalling dynamics were illustrated using Sankey diagrams drawn with pySankey2, and chord diagrams in polar coordinates were rendered using pyCirclize. Finally, H&E-stained tissue sections were processed and visualised with SpatialData [32], enabling spatially resolved integration of histological and transcriptomic signals.

## Results

### ST resolved at single cell level is suitable for integrated analysis of histology and spatial expression profiling to elucidate the TME

First, we examined the spatial profiles and cellular composition of a Human Lung Cancer sample using Xenium Prime. A representative region of 1 mm^2^ was selected using the highest selection score. The hematoxylin and eosin staining (Fig. S1A) revealed heterogeneous histological structures, with regions of solid tumour growth and dense immune cell infiltration, particularly concentrated in the right area. The spatial distribution of TME constituent cells (Supplementary Fig. S1B) revealed that lymphocytes, including CD8^+^ T cells, infiltrated the tumour parenchyma. Notably, CD8^+^ T cells (in navy) were spatially enriched at the tumour–immune interface, suggesting potential cytotoxic activity. To further probe immune functionality, we visualised the spatial expression of the immune checkpoint genes *PDCD1* (encoding PD-1) and *CTLA4*. Both the genes exhibited spatially restricted expression patterns (Supplementary Fig. S1D– E), with high levels near the tumour margins, suggesting locally activated or exhausted T-cell phenotypes. These findings provide a high-resolution view of not only the histomorphology but also the spatial expression landscape, facilitating TME characterization.

This pattern was also confirmed by Xenium Prime data from Human Breast Cancer (Supplementary Fig. S2). As expected, immune infiltration within the tumour parenchyma was observed in the representative region, accompanied by low expression of checkpoint genes.

### GraphTME elucidates individual interactions between cells within the TME

We extended beyond these traditional phenomenological approaches to elucidate and visualise spatial interactions and functional relationships between all cells within the region. Interactions within the representative region of the MERFISH dataset from patient Pt73, reported by Chen et al., are shown in Fig. 2. We stratified the inferred interactions by signalling pathway to examine functional differences in cell–cell communication within the tumour microenvironment (Supplementary Fig. S3). In the checkpoint and growth factor pathways, tumour cells and lymphocytes contributed comparably. In contrast, the cytokine pathway exhibited prominent and targeted signalling from T cells to antigen-presenting cells—such as macrophages and conventional dendritic cells type 2 (cDC2)—indicating a focused immune activation.

**Fig. 2:**
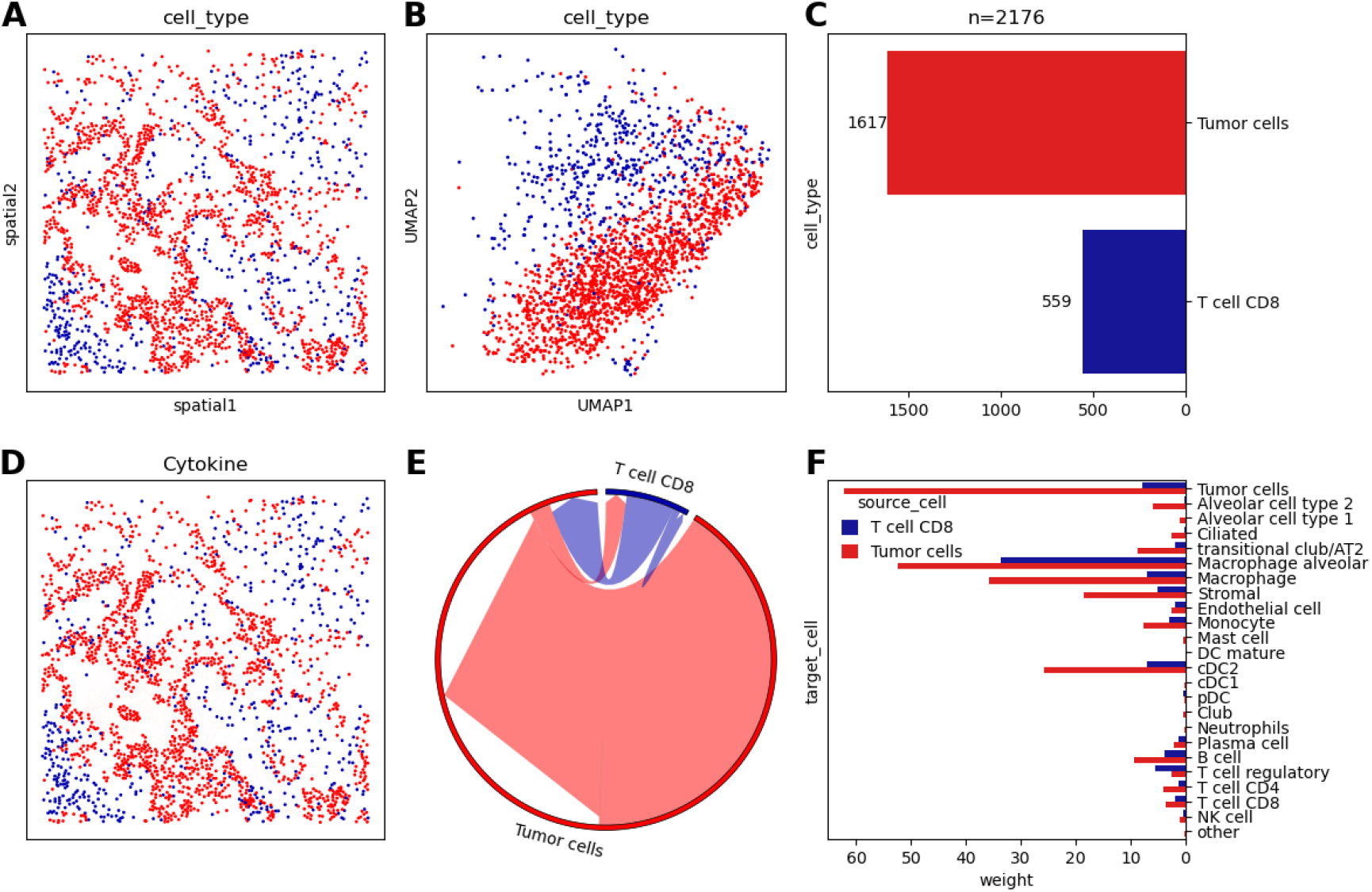
Results of CCI within the representative region of MERFISH for an NSCLC sample. **A**. Spatial distribution of tumour and CD8^+^ T cells; **B**. Projection of gene expression onto the embedding space; **C**. Cell counts for each cell type; **D**. Interaction edges between cells in the cytokine signalling pathway; edge thickness is normalised, and edge colours correspond to the source cell type; **E**. Chord diagram showing aggregating interaction weights between each pair of cell types; **F**. Aggregated outgoing interaction weighs toward other cells, grouped by source cell type.

This suggests that cytokine signalling plays a central role in immune activation, with T cells engaging immune effectors in a focused and coordinated manner.

We next focused specifically on interactions between tumour cells and CD8^+^ T cells, as they are key to immune phenotype characterization. From the spatial distribution in Fig. 2A, CD8^+^ T cell infiltration into tumour tissue was evident. The corresponding gene expression patterns projected onto the 2D tissue surface (Fig. 2B) revealed distinct expression profiles for each cell type. Supplementary Fig. S4 shows the aggregated pathway weights highlighting functional interactions between these two cells. Among the enriched pathways, we selected the cytokine-mediated signalling pathway as representative for analysing the immune response. Fig. 2D visualises individual cell-dell interactions as edges, highlighting spatial regions where communication is most prominent. A chord diagram summarising the bidirectional communication between CD8^+^ T and tumour cells is shown in Fig. 2E. The cumulative interaction weights confirmed that cytokine secretion from CD8^+^ T cells to tumour cells was disproportionately high, despite the numerical inferiority of CD8^+^ T cells (Fig. 2C). Finally, Fig. 2F shows the total cytokine signalling output from CD8^+^ T cells and tumour cells to all other cell types. Both cell types interact with a diverse set of targets, including macrophages, stromal cells, endothelial cells and lymphocytes. Here, in lung cancer tissue, macrophage alveolar cells are often considered phenotypically similar to tumour cells. More specifically, the CD8^+^ T cell signalling was directed toward major antigen-presenting cell types and other immune populations. These findings suggest that cytokine signalling within the TME contributes to active immunomodulation through cell-type-specific communication strategies.

### GraphTME model predicts ICI response for NSCLC with high accuracy

We not only examined the interaction patterns of cells but also built a model to predict responsiveness to the ICI treatment, using Chromium-based single-cell RNA-seq data from CD8^+^ T cells of NSCLC patients treated with ICIs. Substantial inter-patient variation was observed prior to correction (Fig. 3A), which was reduced after batch correction (Fig. 3B). Clinical annotations based on binary anti-PD-1 response labels (Fig. 3E) were used to supervise model training. We confirmed that the expression patterns of responder and non-responder cells separated in a 2D latent space. To ensure class balance, the numbers of responder and non-responder cells were equalised as far as possible (Fig. 3F). We next examined differential gene expression between predicted responders and non-responders (Fig. 3G). Responder cells expressed higher levels of *CCL5*, a prognostic biomarker for immunotherapy [33], whereas non-responders showed elevated expression of *LTB*, a marker associated with cancer progression and metastasis [34]. The dataset was divided into training (P1, P2, P3, P6, P7-pre) and test (P4, P5, P7-post) sets, as shown in Fig. 3H. This split ensured that more cells were used for training than for testing (Fig. 3I), with no patient-level overlap between sets. Evaluation results of the ICI response prediction model and confusion matrices (Fig. 3J and 3K), showed high sensitivity and specificity. We also confirmed that the *F*_1_ score exceeded 0.83. These results indicate that the GraphTME model generalises well across patients and accurately captures clinically relevant immune phenotypes for predicting ICI responses from spatial CD8^+^ T cells.

**Fig. 3:**
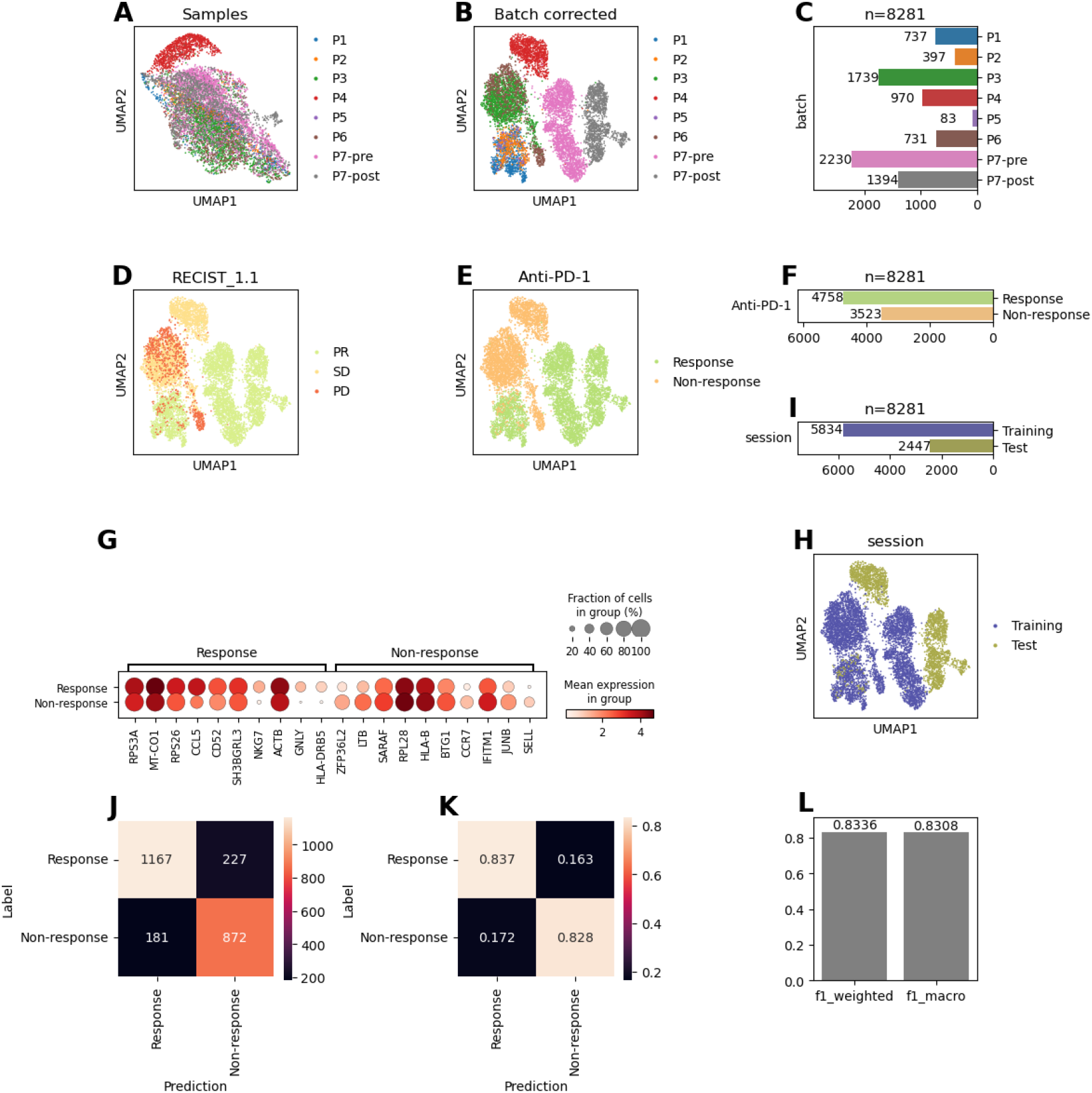
Training and evaluation of the ICI prediction model using Chromium-based CD8^+^ T cell data from patients with NSCLC with anti-PD-1 responses. **A**. 2D projection of CD8^+^ T cells coloured by patient identity (P1–P7). ; **B**. 2D projection after batch correction; **C**. Number of CD8^+^ T cells per patient; **D**. 2D projection coloured by clinical response based on RECIST 1.1; **E**. 2D projection coloured by binarised anti-PD-1 response labels; **F**. Number of responder and non-responder cells; **G**. Dot plot of differentially expressed genes between responders and non-responders; **H**. 2D projection split into training and test sets for model evaluation; **I**. Number of cells used for training and testing; **J**. Confusion matrix model predictions on the test set; **K**. Row-normalised version of the confusion matrix; **L**. *F*_1_ scores on the test set.

### The result of ICI response prediction by the GraphTME model was validated to be identical to the actual results

To assess the predictive capacity of the GraphTME model on actual ST data, we extracted CD8^+^ T cells from the MERFISH dataset on NSCLC patient cases and performed inference on these at the single-cell level. The results are shown in Fig. 4. As illustrated in Fig. 4A, the predicted responder and non-responder cells were spatially intermingled. In contrast, the 2D projection of the model’s learned embeddings (Fig. 4B), which integrate both node and edge features, revealed a clear separation between response states. This indicates that the latent encoding based on functional differences effectively captures biologically meaningful variation and serves as a foundation for response classification. As a demographic aggregation between the two groups, a comparison of the number of cells in Fig. 4C showed that most cells were predicted to be responders, which was consistent with the patient’s known clinical PR status. To gain mechanistic insight, we examined differential gene expression between predicted responders and non-responders in Fig. 4D. The *ANXA1* gene, which is prominent in the responder group, is known as a potential anticancer agent that inhibits the NF-*κ*B signalling pathway [35]. In contrast, the *COL3A1* gene, which was prominent in the non-responder group, is known to be associated with poor prognosis, such as tumour growth and metastasis in NSCLC [36]. In addition, Fig. 4E shows the weighted sum of target cells interacting with the two cell groups. Responder cells directed more total signalling toward tumour cells and macrophages. This indicates that the immune-related signalling of the responder cells was more dominant. These findings suggest that spatially localised CD8^+^ T cells with effector-like signatures are functionally involved in antitumour responses.

**Fig. 4:**
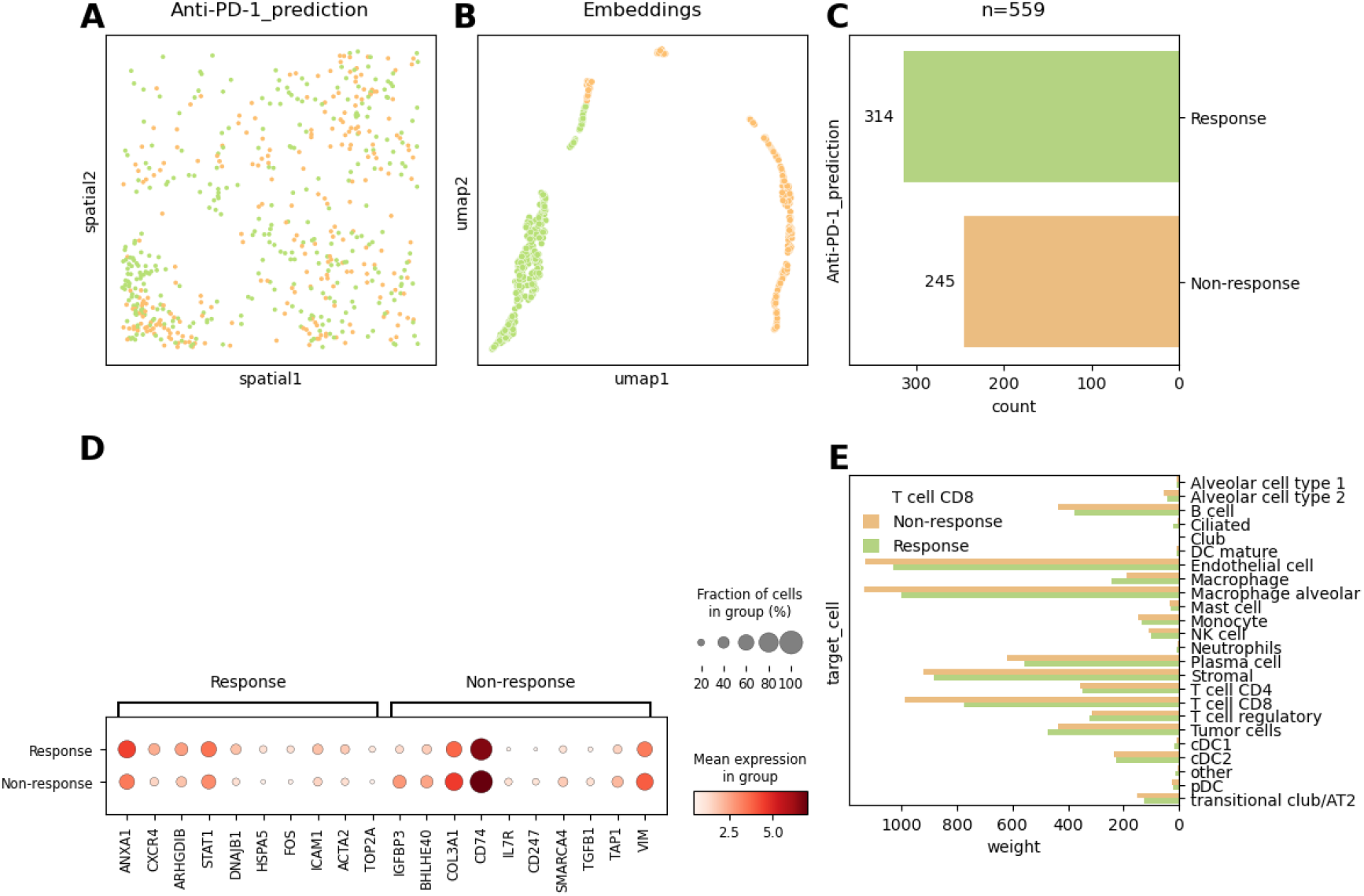
Single-cell-level prediction of anti-PD-1 response for CD8^+^ T cells within the representative region of MERFISH for an NSCLC sample. **A**. Spatial distribution of CD8^+^ T cells classified as responders or non-responders; **B**. 2D projection of the model-derived embedding vectors coloured by predicted response class; **C**. Cell count of predicted responders and non-responders; **D**. Dot plot of differentially expressed genes between predicted responders and non-responders.; **E**. Aggregated signalling weights from CD8^+^ T cells stratified by predicted ICI response toward all other cell types.

## Discussion

Understanding how tumour-infiltrating immune cells interact with the surrounding cells within the spatial organisation of the TME is critical for advancing precision immunotherapy. In this study, we introduced GraphTME, a spatially grounded, biologically informed, graph-based modelling framework that predicts ICI responses at single-cell resolution by explicitly modelling pathway-specific CCIs. Our approach leverages curated ligand-receptor interactions, spatial proximity, and transcriptomic signatures to construct multirelational graphs and applies RGCNs for predictive modelling. To the best of our knowledge, GraphTME is the first framework that links spatially resolved CCIs to functional immunotherapy response inference *in situ*.

While existing methods are limited to the descriptive modelling of CCI, GraphTME has three distinct points. First, it analyses ligand-receptor interactions via signalling pathways so that they can be interpreted in the context of immune surveillance and tumour evasion. Second, to our knowledge, this is the first framework that uses spatially resolved interaction graphs between individual cells to predict the anti-PD-1 response *in situ*. Third, it provides clear evidence and quantitative outputs, rather than just trends or patterns. Thus, GraphTME is a robust and interpretable tool for both discovery and diagnosis.

This GraphTME model is also flexible in terms of scalability; it is not limited to the ST platform but is more suitable for imaging-based ST, which is advantageous for cell-level studies, than sequencing-based ST, which is advantageous for whole-transcriptome analysis. In addition to the lung and breast cancers presented in this study, it can be applied to other cancer types, provided that appropriate datasets are available.

Despite these promising results, this study had several limitations. First, publicly available datasets of clinical ICI outcomes are scarce. Prior studies [37, 38, 39] have relied on bulk expression profiles, limiting their utility for fine-grained single-cell-level modelling. To overcome this limitation, larger prospective cohorts are required. Second, although we used the TICCom database to derive pathway-specific ligand-receptor interactions, many entries were not annotated with pathway information, making interpretation challenging. Efforts to curate and expand these resources will improve the interpretability of CCI graphs. Third, only the representative region where CD8^+^ cells infiltrated the tumour tissue was selected as the region of interest, owing to the limitations of the interaction operation. Even with vector operations and parallel processing, there are limitations in processing edges, which can reach hundreds of millions. However, because the selection score was quantified and re-examined in terms of cytomorphology, it was validated for use as a representative region to assess ICI responsiveness. A previous study examining the correlation between CD8+ T-cell density on immunohistochemistry images and survival in patients with prostate cancer also observed a region of 1 mm^2^ [40]. In addition, a 1 mm^2^ patch was used in a previous study that classified the immune phenotype of cytotoxic T cells from H&E images of NSCLC patients [41]. Similar to ST, recent studies have explored transcriptome-based graph neural network (GNN) approaches to predict ICI responses, particularly using bulk expression data and pathway-level features. Among these, ICInet [42] and IRnet [43] represent applications of graph learning to transcriptomic data integrated with prior biological knowledge. Although the target cells were not specified and spatial resolution was lacking, their emphasis on pathway-driven modelling aligns conceptually with our proposed framework. ICInet demonstrates the strength of leveraging curated gene-gene interaction networks and pathway knowledge to guide feature selection. IRnet, on the other hand, emphasises interpretability by projecting gene expression onto pathway embeddings and learning inter-pathway relationships via attention-based GNNs. These studies underscore the suitability of GNNs for modelling immune responses from transcriptomic data while emphasising the importance of incorporating pathway-aware biological contexts to enhance interpretability and predictive performance.

The GraphTME framework offers a promising approach for clinical applications. This may serve as a predictive biomarker for selecting patients who are most likely to benefit from anti-PD-1 therapy. In immunotherapy, which has a low response rate, the use of this prediction model is expected to reduce unnecessary treatment burden [44]. It can also be used for the quantitative comparison of the TME before and after immunotherapy by tracking changes in spatially resolved CCI patterns, immune cell infiltration, and pathway-specific signalling strength. This framework provides a future direction for research on cellular interactions in the context of spatial biology. Bridging microscopic cellular reactions to macroscale clinical phenotypes may be a milestone in the field of translational medicine. As ST technologies become more widespread, modelling the spatial logic of CCIs will be useful not only for immuno-oncology, but also for neurodegeneration and infectious disease research [45].

## Conclusion

In summary, GraphTME bridges the gap between descriptive ST and functional immunotherapy modelling by enabling interpretable, single-cell-level prediction of ICI responses based on spatially resolved CCIs. This integrative framework may serve as a foundation for future efforts to personalise cancer immunotherapy using spatially grounded computational models.

## Supplementary data

Supplementary data is available in online.

## Author contributions statement

Conceptualisation: H.J.; Data curation: H.J.; Formal analysis: H.J.; Investigation: H.J.; Methodology: H.J.; Project Administration: H.J.; Sources: H.J. and Y.C.; Software: H.J.; Supervision: H.J.; Validation:H.J., J.O., and Y.C.; Visualization: H.J.; Writing the original draft: H.J.; Writing, review, and editing: All authors.

## Acknowledgments

The authors thank the anonymous reviewers for their valuable suggestions. The authors used ChatGPT, a generative artificial intelligence service, to refine English phrasing and conduct minor code reviews. The authors reviewed and edited the content and take full responsibility for it.

## Data Availability

Single-cell reference data were obtained from the CELLxGENE Discover datasets: https://cellxgene.cziscience.com/datasets. Tumour-immune system-specific ligand-receptor interactions were derived from the following TICCom: https://github.com/yunjinxie/TICCom-dataset. Public data on cancer samples from Xenium Prime platforms were obtained from the 10X Genomics datasets page: https://www.10xgenomics.com/datasets/.

## Code Availablity

The code for the GraphTME model is maintained in the following GitHub repository: https://github.com/recognizability/graphtme

---

**Supplementary Fig. S1:**
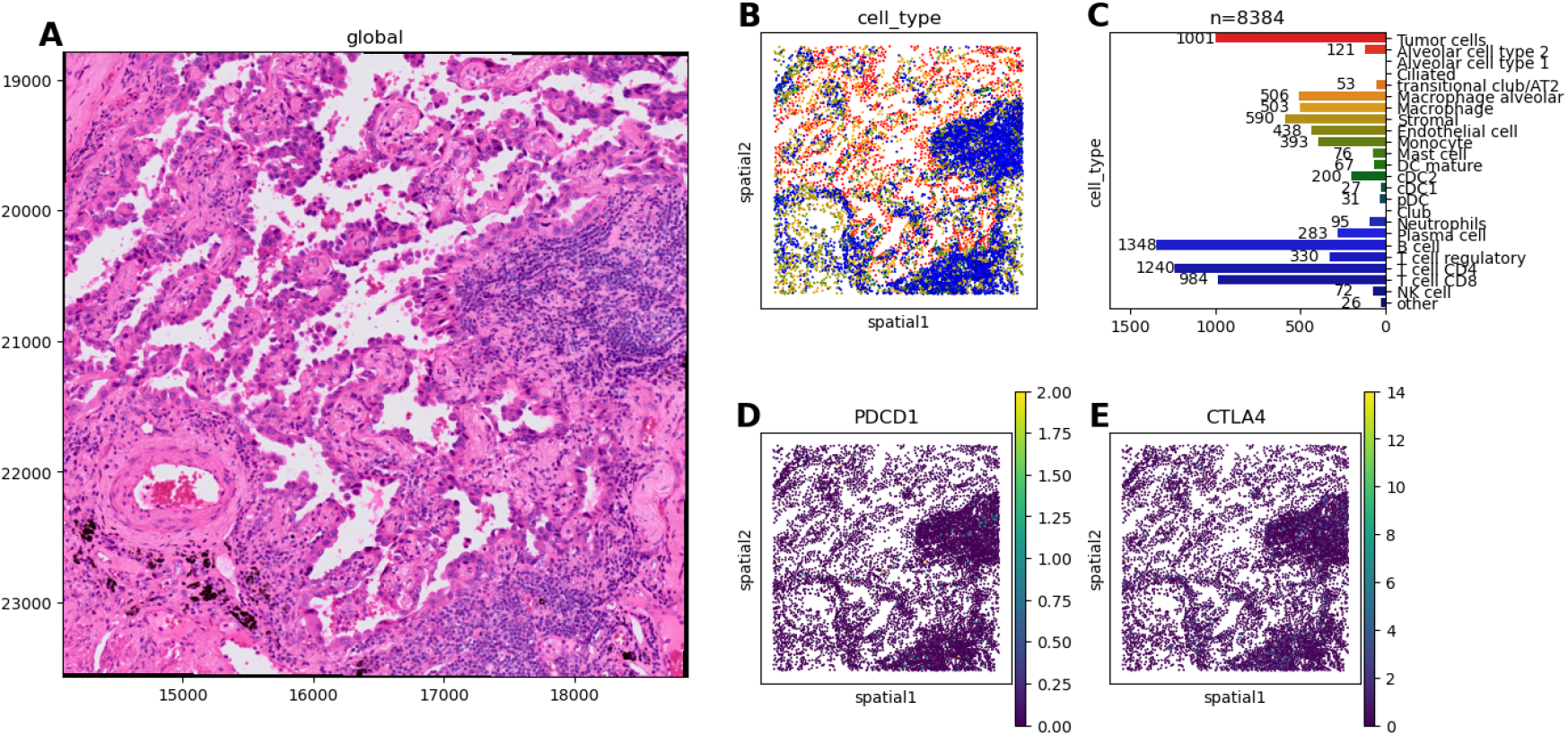
Histological and spatial transcriptomic profiles on the same coordinate system within a representative region of Xenium Prime from a Human Lung Cancer sample. **A**. H&E-stained histological image of a 1 mm^2^ area where immune cells have infiltrated the tumour in the right area; **B**. Spatial distribution of cell types that constitute the TME; **C**. Number of cells of each type; **D-E**. Spatial expression of key immune checkpoint genes, *PDCD1* (encoding PD-1, **D**) and *CTLA4* (**E**), plotted on a per-cell basis. Expression is concentrated in the immune-rich region, especially near tumour boundaries.

**Supplementary Fig. S2:**
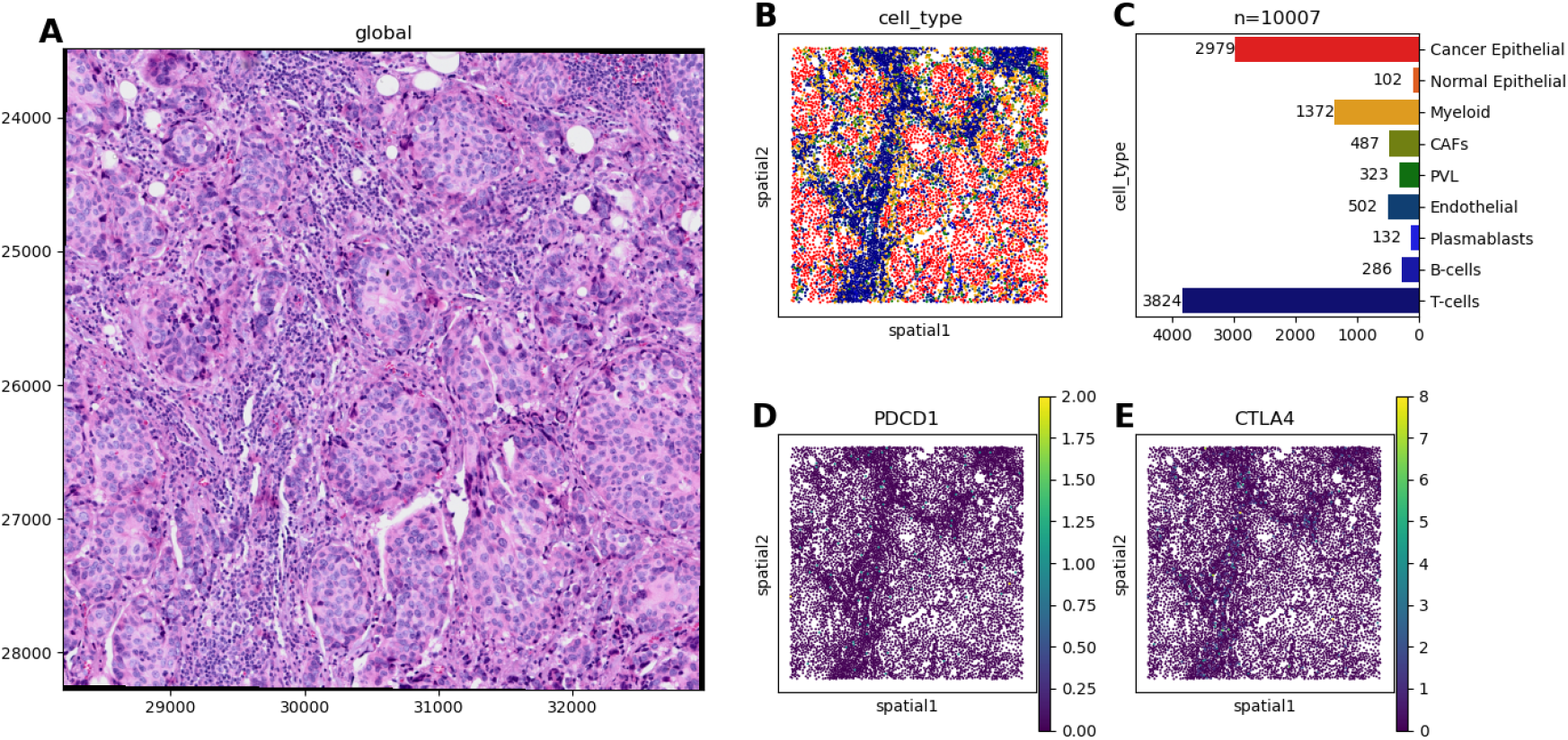
Histological and spatial transcriptomic profiles on the same coordinate system within a representative region of Xenium Prime from a Human Breast Cancer sample. **A**. H&E image of a 1 mm^2^ area where immune cells have infiltrated the tumour area; **B**. Spatial distribution of cell types that constitute the TME; **C**. Number of cells of each type; **D–E**. Spatial expression of key immune checkpoint genes *PDCD1* (**D**) and *CTLA4* (**E**). Expression is scattered and primarily localised to T cell-rich areas.

**Supplementary Fig. S3:**
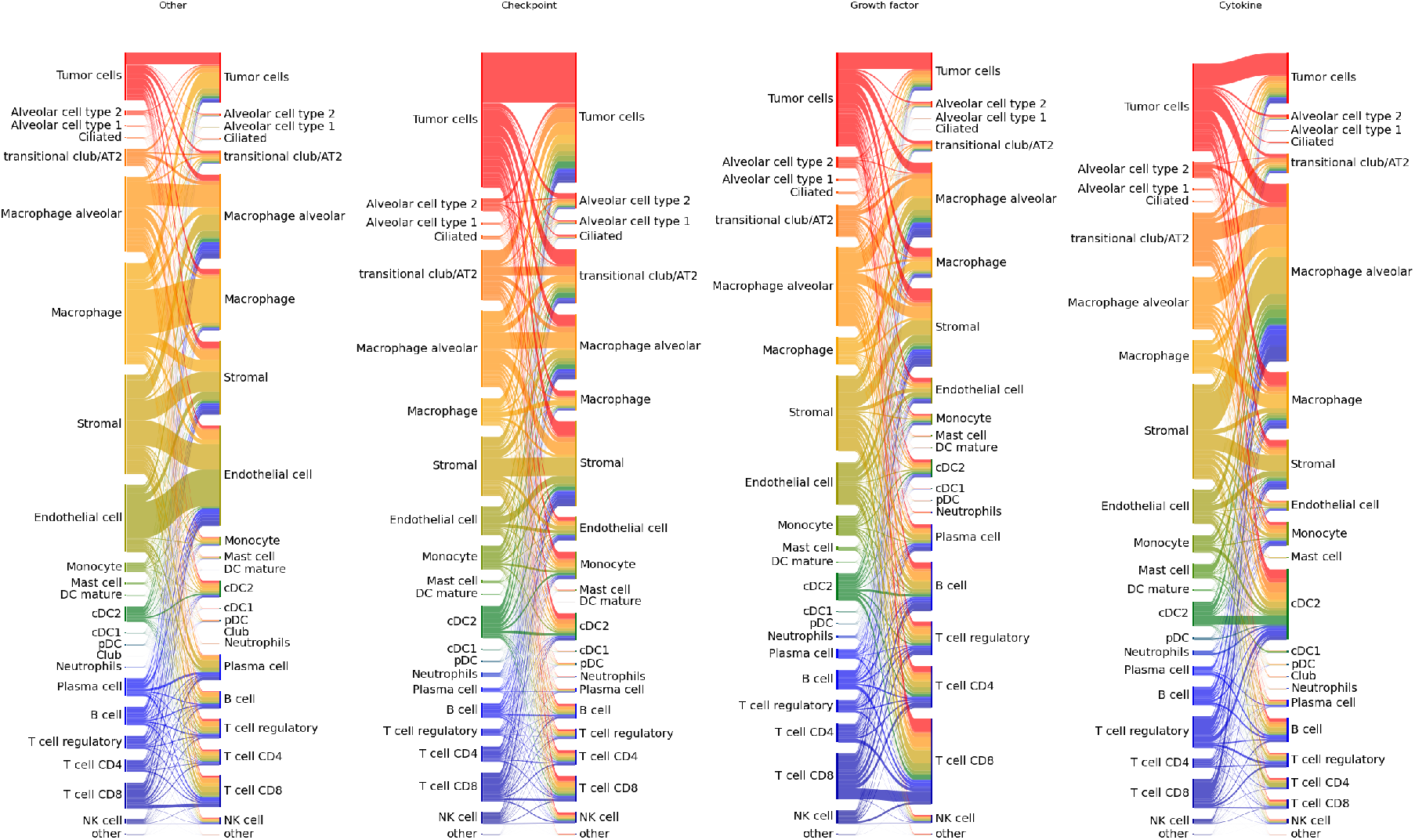
Pathway-specific cell–cell interactions within the tumour microenvironment for representative regions in MERFISH data from a NSCLC case. Sankey diagrams depict directed interactions between all cell types across four signalling pathway groups (Other, Checkpoint, Growth factor, Cytokine). Edge width reflects the cumulative interaction strength for each ligand–receptor pair within the specified pathway category.

**Supplementary Fig. S4:**
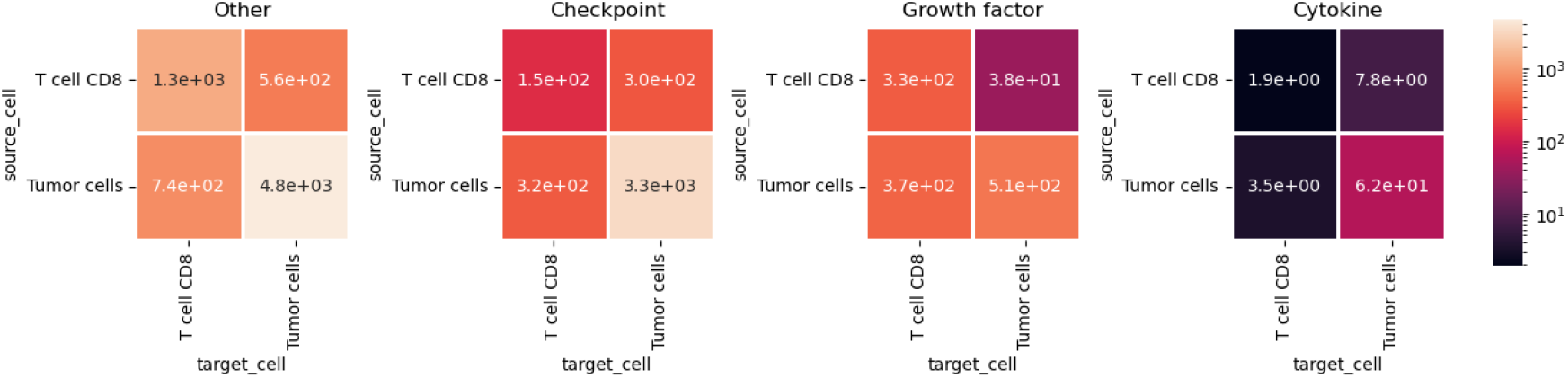
Summed interaction weights between the tumour cells and the CD8^+^ T cells for each of the four pathways for representative regions in MERFISH data from a NSCLC case. Pathways are ordered by total weights, and colours are shown on a log scale.

